# Asynchronous temporal variance in learning behaviour and neural gene expression in a butterfly

**DOI:** 10.1101/2024.03.07.583937

**Authors:** Yi Ting Ter, Erica L. Westerman

## Abstract

Mate preference learning, including imprinting-like learning, is pervasive across animal taxa, and can affect the selection and maintenance of certain phenotypes. However, not much is known about the temporal dynamics behind imprinting-like learning, or the genetic underpinnings underlying it. To uncover the temporal dynamics of imprinting-like learning, from both a behavioural and transcriptional perspective, we conducted learning and RNA-Seq time series using the butterfly *Bicyclus anynana*, a species where both sexes learn mate preferences. We exposed females to an unfamiliar, unpreferred male phenotype (4-spotted male) for five different exposure (training) periods, ranging from 30 minutes to three hours, and recorded their choice between the preferred, familiar male phenotype (2-spotted) and 4-spotted males in mate choice trials conducted two days after training. We also assessed differential gene expression of naïve females and females exposed to trainer males for these same five exposure periods, to identify temporal patterns in gene expression associated with learning. While we found that the longest exposure had the strongest effect on preference learning, we did not observe a linear effect of exposure time on learning. We also show that the highest peak of differentially expressed genes (DEGs) was after one hour of exposure. While a number of genes were uniquely DE at each time point, one gene, associated with transcription initiation, was differentially expressed during learning across all five time points. We observed a similar decreasing trend in both gene expression and learned response after 1.5 hours of exposure, offering new insights to possible attention and forgetting mechanisms. Therefore, our results indicate that both gene expression and learning are temporally dynamic, and not linear through time. They also highlight the role of transcription in mate preference learning and memory formation in Lepidoptera, and illustrate that imprinting-like learning may exhibit similar molecular temporal dynamics as associative learning.

## Introduction

Learning is an important process in the survival and reproduction of animals [1,2]. An animal that changes its behaviour because of an experience is said to have learned, and therefore able to behaviourally adapt to the new environment [2,3]. This change in behaviour is associated with experience-dependent plasticity in the central nervous system, which allows an animal to adapt its responses to stimuli over different time scales [4,5]. For example, an animal that spends a few seconds with a potential mate might make a less informed mating decision compared to one who spends minutes or hours with a potential mate [6–8]. In nature, animals have limited amounts of time to acquire relevant information, learn, and modify their behaviour in response to stimuli. Therefore, the length of exposure an animal has to a stimulus might affect whether, or how much, novel information is retained and cemented into memory. This has been reflected in studies examining training time, learning and memory in animals in a variety of contexts [9–13], where individuals become better at learning the longer they are exposed to a stimulus, indicating that length of exposure positively influences memory formation. While the mechanisms behind learning and memory formation have been explored in both vertebrates and invertebrates, most studies to date focus on associative tasks, like associating colour or odour to a food reward/ punishment [3,9,11–13]. Relatively little is known about the temporal dynamics involved in imprinting-like learning (the early exposure of sexually immature adults to other adult phenotypes [3,14–16]), much less the genetic underpinnings behind this type of learning, especially in the context of sexual ornaments.

The genetic underpinnings of associative learning and memory formation, however, have been well studied in model animal taxa, and include genes associated with the regulation of transcription and translation, hormones, sensory receptors, and neurotransmitters. For example, genes that encode for transcription factors, known as immediate early genes (IEGs; e.g. *egr1, c-fos, c-jun*) are widely used as molecular markers for synaptic plasticity underlying long-term memory formation due to their transient and rapid expression after stimulation [17–19]. Hormones have also been implicated in memory formation, such as 20-hydroecdysone (20E; [20]) and juvenile hormone (JH; [21]). These studies have identified two time sensitive windows for long term memory formation, with the first window being when IEGs are expressed rapidly in response to a stimulus (around initiation of training to one hour later), while the second window is when late response genes (i.e. genes that are expressed later in response to previous signalling from IEGs) were expressed (roughly three to eight hours later) [22–27]. However, most mechanistic studies to date have been conducted on the learning of simple associative tasks, relatively little is known about the temporal genetic architecture underlying imprinting-like learning, or whether imprinting-like learning follows similar transcriptomic temporal patterns as associative learning. Since imprinting-like learning is often hypothesized to be cognitively different from associative learning, due to the implied lasting effect of the training and the higher difficulty of reverse learning [28–30], illuminating its temporal dynamics may give us new insights into the mechanistic similarities (and differences) between associative and imprinting-like learning. In addition, to our knowledge, no studies have looked at the temporal dynamics of learning behaviour and its associated gene expression together, usually these are assessed independently, either in systems conducive to learning time series, or systems conducive to transcriptomic series [22,26,27].

To address these gaps in literature, we used the butterfly *Bicyclus anynana* to examine the temporal dynamics behind imprinting-like learning and its associated gene expression through increasing exposure (training) durations. Female and male *B. anynana* butterflies learn mate preferences for visual signals if they are exposed to sexually mature individuals of the opposite sex shortly after emergence from chrysalis, a time when neither males nor females are ready to mate in this species [31]. Specific to females, naïve females prefer males with two dorsal forewing eyespots, however females exposed to males with four dorsal forewing eyespots immediately following emergence prefer four spot males over two spot males in later mating trials [16,32–34]. Past studies using *B. anynana* to investigate visual imprinting-like learning, as well as the associated brain gene expression, have typically used three hours as the exposure period. However, it remains unclear whether three hours of exposure is the optimal time point for behavioural and gene expression analyses in this study system, or butterflies in general, as no time courses for either have been performed. It is also beneficial to understand whether, or how, duration of exposure time matters, both for downstream behavioural responses and for identifying genes and gene networks associated with the mate preference learning process. In an ecological context, it is likely that animals are exposed to other conspecifics and heterospecifics for a range of duration times, and this variation in social exposure time may influence the evolutionary importance of learned versus innate mate preferences. In butterflies, some species roost at night or overwinter, therefore interacting with one another for long periods of time [35–37]. Shorter interactions are also likely, and therefore may have the potential to also influence subsequent mating decisions. Neurogenomically, transcriptional changes responsible for long-term memory formation are initiated shortly after exposure in all animal systems studied to date [22,26,27]. Therefore, by only assessing transcriptional change at the end of a three-hour (or any lengthy exposure) period [34], we are likely missing genes critically important to mate preference learning and memory, which would better help us understand the mechanistic similarities between imprinting-like learning and associative learning in general.

Therefore, in this study we examined the effect of increasing exposure time on learning ability by exposing naïve females to 4-spotted males for five different time periods (i.e. 30 minutes, 1 hour, 1.5 hours, 2 hours and 3 hours) and later testing female preference for four spot males relative to that of socially naïve females. Additionally, to examine the effect of exposure time on gene expression associated with learning, we then performed RNA-Seq analyses of female heads collected at these same training time points, which elucidated what genes were differentially expressed at different training time points, and whether amount of gene expression is predictive of extent of future behavioural change. We then performed a weighted gene co-expression network analysis (WGCNA), allowing for temporal visualisation of gene suites and revealing gene co-expression networks associated with mate preference learning.

## Results

We found that the duration of exposure time to 4-spotted males influenced learning a new spot preference in female *B. anynana*. As the duration of exposure time increased from 30 minutes to 3 hours, learning was shown to be dynamic through time. Females learned to prefer 4-spotted males significantly after 3 hours of exposure (p = 0.008; χ^2^ = 6.91) compared to naïve females, but showed no significant change in preference at other time points (Fig 1A). Interestingly, despite an increasing trend (albeit not significant) to prefer 4-spotted males as exposure time increased from 30 mins to 1 hour, there was a decreasing trend in preference for 4-spotted males at the 1.5-hour time point (p=0.08; χ^2^ = 2.88), to similar levels as naïve females (Fig 1A).

**Fig 1.**
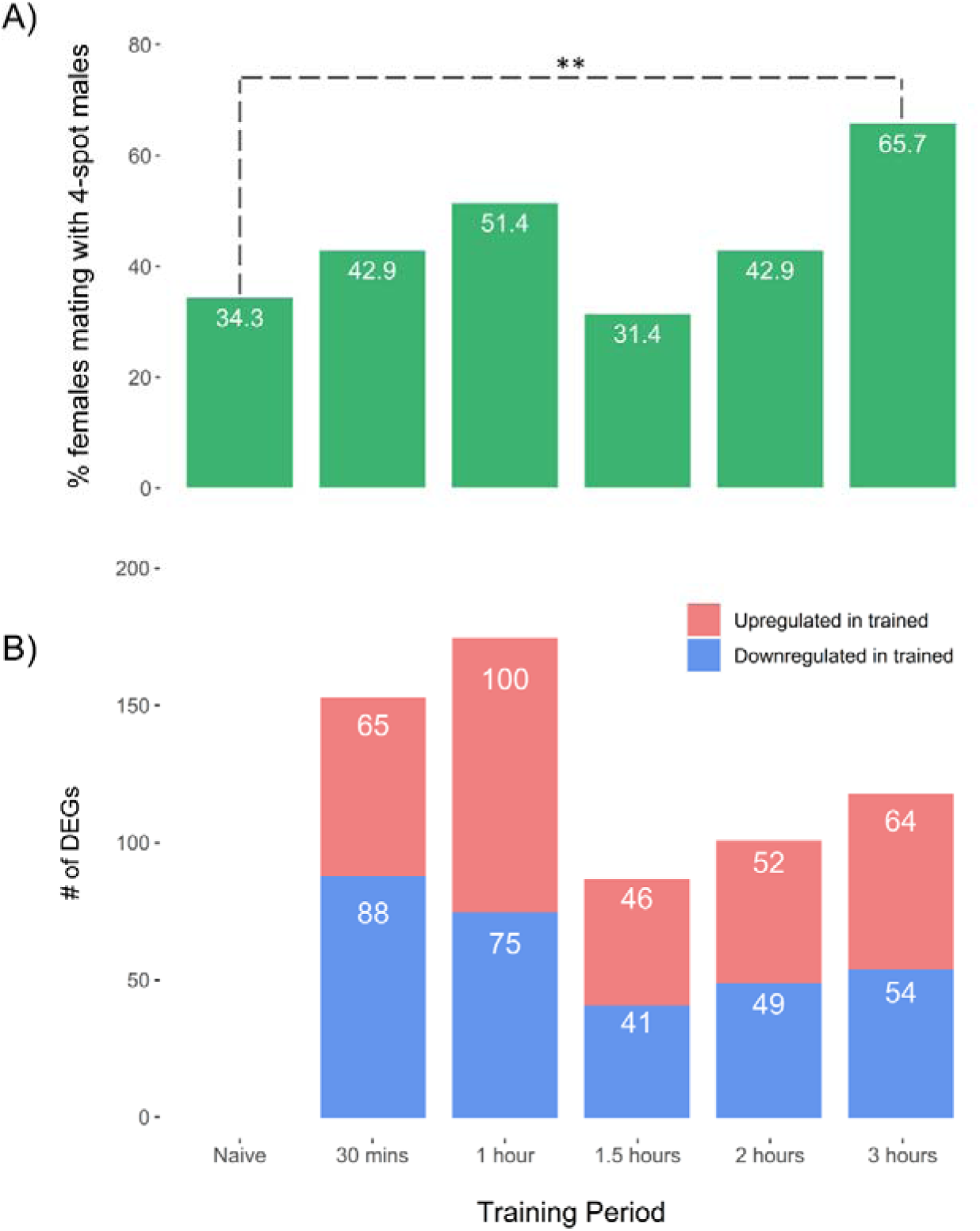
Temporal dynamics of learning and its associated gene expression across five exposure time points – 30 minutes, 1 hour, 1.5 hours, 2 hours and 3 hours of exposure to 4-spotted males. (A) Percentage of females that mated with 4-spotted males post-training across five exposure periods (n = 35 per exposure period). (B) Number of differentially expressed genes (DEGs) between trained and naïve females across five time points (n = 3 per treatment and time point). Blue indicated downregulated genes and red indicated upregulated genes.

To explore gene expression changes that correspond to increasing exposure time, we sequenced whole-head transcriptomes of naïve and trained female *B. anynana* at five different time points – at 30 minutes, 1 hour, 1.5 hours, 2 hours and 3 hours (N = 3 per treatment per time point; N = 30 total). This generated a total of approximately one billion high quality 100 base pair, single-end (SE) reads (Table S1). Adapter trimming removed 0.2 million reads (0.02%) and approximately 90% of the remaining trimmed reads mapped to the *B. anynana* genome v1.2 [60]. Across all libraries, 16,620 genes (73% of 22,642 annotated genes in the genome) had at least 10 mapped reads, and were used for differential expression analyses.

We found that gene expression associated with learning was also temporally dynamic (Fig 1B). The number of differentially expressed genes (DEGs) peaked after 1 hour of exposure (175 DEGs; Figs 1B, 2C, Table S3), and was lowest after 1.5 hours of exposure (87 DEGs; Figs 1B, 2D, Table S4) There was a correlational link between gene expression and learning, in which the 1.5-hour time point was the lowest in both assays. However, there was a difference in peak effects, as peak DEG was after 1 hour of training (Fig 1B), while peak behavioural response was after 3 hours of training (Fig 1A).

**Fig 2.**
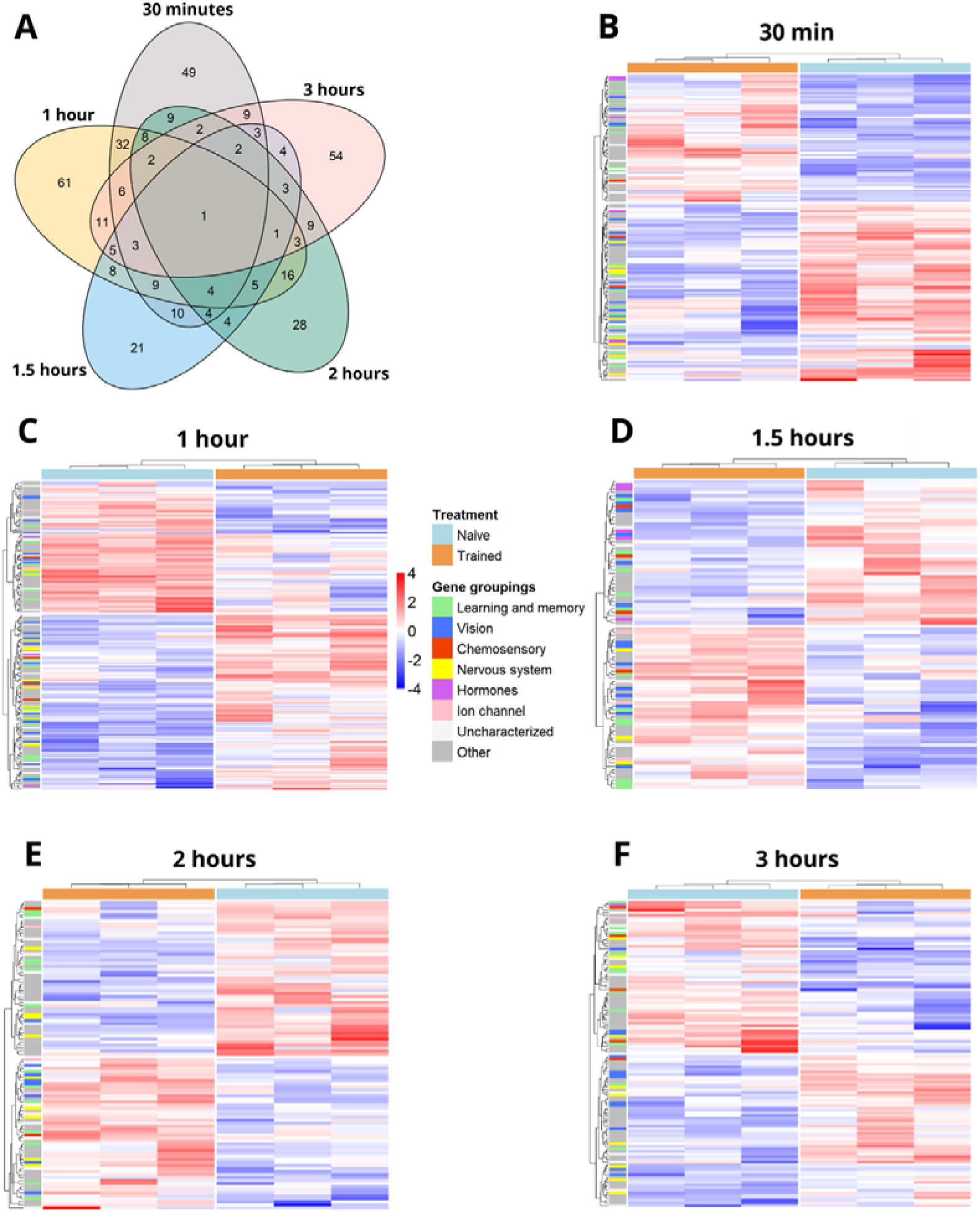
DEGs between trained and naïve females across five time points. **(A)** Venn diagram showing the overlaps for DEG between the five time points. One DEG, transcription initiation factor TFIID subunit 6, is shared across all five time points. **(B-F)** Each row indicates a single gene, and each column indicates a single sample. Counts were normalized using variance stabilizing transformation, and Z-scores were calculated for plotting. Genes and samples were clustered by expression patterns, with blue indicating decreased expression and red indicating increased expression. Each gene was further categorized into a gene group, based on putative function. Heatmap of DEGs between trained and naïve females that were exposed to males for **(A)** 30 minutes; **(B)** 1 hr; **(C)** 1.5 hrs; **(D)** 2 hrs; and **(E)** 3 hrs.

When comparing across time points, we found that there was large overlap of DEGs between the 30-mins and 1-hour time points (25%; 65/263) compared to other time points, with 32 DEGs specifically differentially expressed in these two time points only (Fig 2A). There was also more learning and memory genes and visual genes during the 30-minute and 1-hour time points respectively (Fig S1; Figs 1B, 1C, Tables S8, S9). Additionally, one DEG, encoding for transcription initiation factor TFIID subunit 6, was shared between all five time points (Fig 2A). While number of DEG differed across training time point, “Naïve” and “Trained” females clustered according to treatment for all time points in our analyses (Fig 2B-F).

To investigate gene networks that are associated with social learning experience, we performed a weighted gene co-expression network analysis (WGCNA), visualising the differential expression of genes across increasing exposure time. We identified 16 modules (Fig 3). We focused on modules with significant expression associations at two time points, the 1 hour and 3-hour time points, given that these were the time points with the highest DEGs and learned response respectively. Of these modules, two modules – the grey60 and lightgreen modules – were of note.

**Fig 3:**
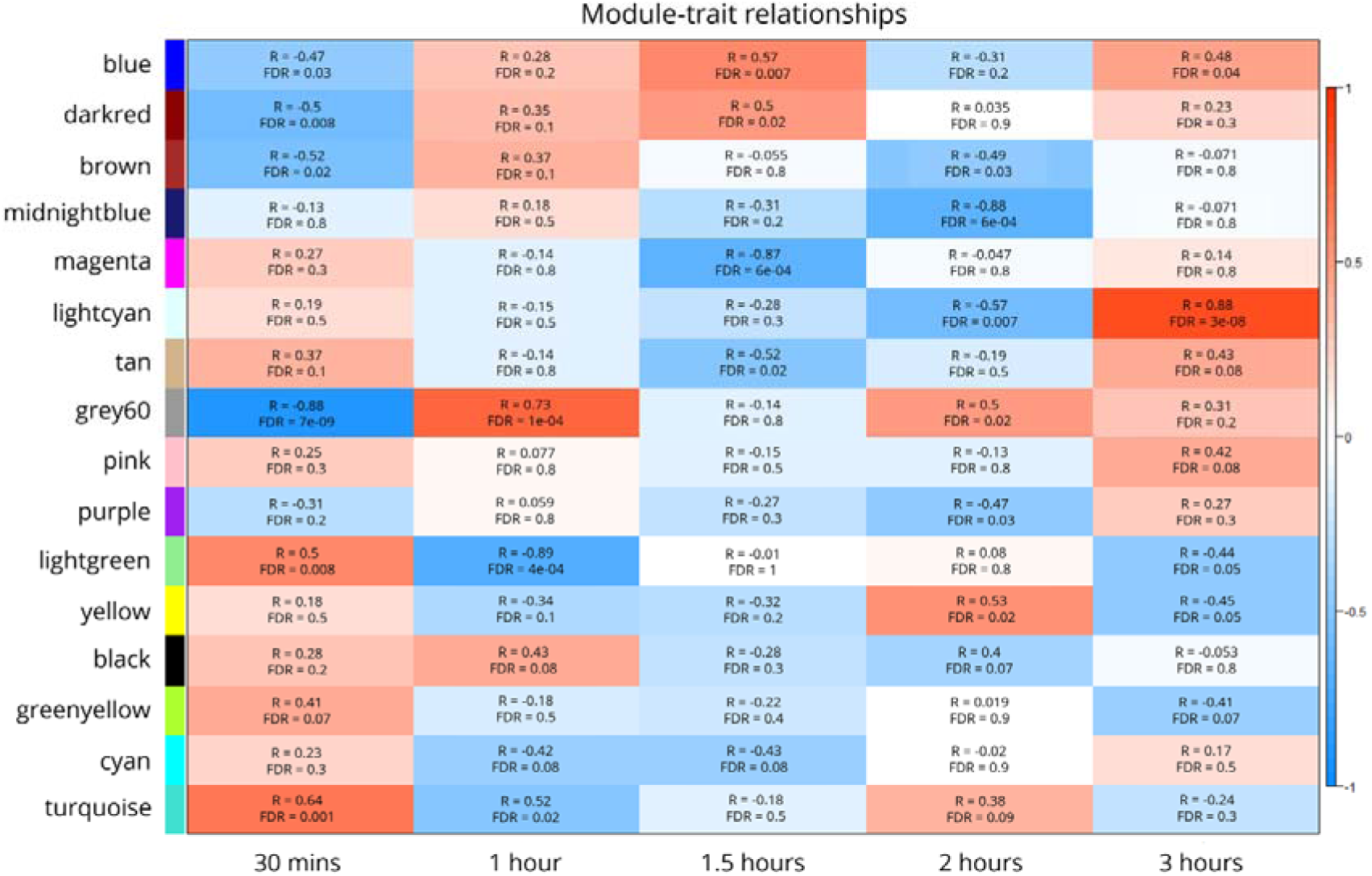
Gene co-expression network modules across five time points. Module-trait association heatmap. Rows indicate module eigengenes (ME) and columns indicate pairwise comparisons between trained and naïve females, across 30 minutes, 1 hour, 1.5 hours, 2 hours and 3 hours. The values in each cell denote the correlation value I and false discovery rate (FDR), with significant FDR values bolded. The intensity of colour corresponds to the strength of association, with r ranging from −1 to 1. Blue indicates lower expression, while red indicates higher expression. Naïve females are set as the reference group.

The grey60 module, consisting of 593 genes, centred around the hub gene uncharacterized protein LOC123868231 (*BANY.1.2.g21105;* Table S17), was significantly downregulated at 30 minutes (r=-0.88, FDR<0.0001), and significantly upregulated at the 1-hour time point (r=0.73, FDR=0.0001). Out of the 593 genes in this module, 51 genes were also identified as DEGs in our study (Table S14). Many of these genes encoded for learning and memory genes, such as autism susceptibility gene 2 protein-like (*BANY.1.2.g6306*), calcitonin gene-related peptide type 1 receptor-like (*BANY.1.2.g01081*), and pumilio homolog 2 (*BANY.1.2.g00387*). However, GO enrichment analysis found no significantly enriched GO terms within the module.

The lightgreen module, consisting of 157 genes centered around the hub gene uncharacterized protein LOC112055531 (*BANY.1.2.g22613*; Table S17), was significantly upregulated at 30 minutes (r = 0.5, FDR = 0.008), significantly downregulated at 1 hour (r = - 0.69, FDR = 0.0004), and significantly downregulated at 3 hours (r = −0.44, FDR = 0.05). Out of the 157 genes in this module, 5 genes were also identified as DEGs in our study (Table S15). GO enrichment analysis found 33 significantly enriched GO terms in the lightgreen module, many of them were involved in tRNA aminoacylation, gene expression, ribonucleotide/ nucleotide binding, ATP binding and metabolic processes (Fig S3, Table S16).

## Discussion

In this study, we showed that the length of exposure to an unfamiliar, sexually mature 4-spotted male affected whether females changed their spot preferences, i.e. whether they learned to prefer 4-spotted males over the innately preferred 2-spot male phenotype in later mate choice assays. We found that peak differential gene expression did not coincide with optimal training time, as peak differential gene expression was after 1 hour of exposure, while optimal training time was after 3 hours of exposure. We also showed that gene expression associated with imprinting-like learning is temporally dynamic, and identified several genes that are differentially expressed during a social learning event. Additionally, one DEG, transcription initiation factor TFIID subunit 6 (*Taf6*), was differentially expressed in the head throughout the entire training process (DE at every time point).

Our result that exposed individuals did not have a continuous increase in learning as exposure time increases was unexpected, as multiple studies using repetitive spaced trials (therefore increasing total exposure time) show a linear relationship between exposure time and learning ability [9–13]. Instead, we showed that there was a decrease in learning from the 1-hour to 1.5-hour time point, in which the percentage of females who learned was similar to naïve preference levels (Fig 1A). It is possible that this decrease in learning could be due to forgetting, which plays an important role in learning and memory [61–64]. Alternatively, this decrease in learning could be the result of changes in male behaviour throughout the exposure period (i.e. males being less active at 1.5 hours). However, prior studies have shown that there is no effect of male behaviour on female preference or female’s likelihood to learn in this system, so it is unlikely that changes in male behaviour during the training period are responsible for the observed effect [16,33,65]. Additionally, since there was a significant increase in females who learned only after three hours of exposure to the training stimulus, this indicates that three hours of exposure is the minimum amount of time required for a female *B. anynana* to learn a new male spot preference. Interestingly, in *B. anynana*, females learn to prefer males with novel enhanced pheromone blends after a 3 min exposure [66]. This shows that learning olfactory cues may require less time than learning visual cues, suggesting that odour may have a higher valence than visual cues. It would be interesting to see if this also holds true in other insects and learning contexts. In *Schizocosa uetzi* spiders, subadult females are also able to learn a male phenotype that they are exposed to. The exposure duration is variable and shorter than our study, with training done every day or alternate day, resulting in at least 30 – 60 minutes of total exposure, but spaced out over time [15,67]. It would be interesting to see if one single long exposure (i.e. 3 hours) would also result in learning in other arthropods instead of multiple learning trials across multiple days.

In this study, we show that the highest number of DEGs was at the 1-hour exposure time point (Fig 1B). This is discordant with the learning curve, in which the highest proportion of females who learned was at the 3-hour exposure time point. Therefore, differential gene expression levels during a training experience are not always predictive of degree of subsequent behavioural response to stimulus, at least in the context of imprinting-like learning. Previous studies have shown that memory consolidation is a temporal process that is dependent on translation and transcription [22–27], and that there are two sensitive time windows related to memory formation, with the first window around training – 1 hour after initial stimulus, and the second window around 3 – 8 hours after initial stimulus [22–27]. Our findings suggest that in *B. anynana*, the first time window of gene expression occurs after 1 hour of exposure. This is likely the phase where immediate early genes (IEGs) are highly expressed, corresponding to the sensitive period of learning and memory in other research [22–27]. However, the second transcriptional time window, suggested to be when genes related to the *de novo* protein synthesis critical for long-term memory formation are expressed, was likely not captured fully in this study [22,27,68,69]. We stopped training, and collecting gene expression data, from females after the 3-hour time point, since that is the training time point that is widely used in other *B. anynana* research and females have successfully learned after that length of exposure to an unpreferred male [16,32]. Future studies could examine gene expression and learning ability past the 3-hour time point, which could reveal genes related to the *de novo* protein synthesis critical for long-term memory formation in Lepidoptera. Since we potentially captured the first window of gene expression, this is likely the time window when immediate early genes (IEGs) are expressed. IEGs are transiently and rapidly expressed in response to a stimulus, before any new proteins are synthesised [18,68–71].

Therefore, neuronal IEGs can be used as an indirect marker to measure neuronal activity, memory formation and other behavioural activities. In our study, we found differential expression of an IEG, the hormone receptor 38 (*Hr38*), which is the insect ortholog of the mammalian Nr4a family of immediate early response transcription factors which have been implicated in long-term memory [17,72–76]. *Hr38* is the first conserved neural activity marker identified in insects, where it has been shown have increased expression in the mushroom body and antennal lobes of male silkworm moth *Bombyx mori* and *D. melanogaster* when exposed to female sex pheromones [73]. Researchers also observed an increase in *Hr38* expression after a 30 min stimulation, with a peak expression at 90 min [73]. This temporal expression pattern corresponds with the *Hr38* expression pattern observed in our study, in which *Hr38* was similarly differentially upregulated in the trained females at the 1.5-hour (90 min) time point (Fig S2). Hr38 expression was also shown to be increased in the mushroom bodies of forager bees compared to nurse bees or queens [77], suggesting a role in spatial learning. A transcriptomic study assessing memory acquisition in *D. melanogaster* also found an upregulation of *Hr38* in the mushroom body, a major site for learning and memory in insects, compared to whole heads [51]. Other transcriptomic studies examining learning and memory across species also recover differential expression of *Hr38/ Nr4a* in their treatments[40,49]. Taken together, these findings strongly imply *Hr38*’s function in learning and memory across learning context in insects, and its being a pertinent candidate gene for future functional studies of imprinting-like learning.

In this study, we studied the temporal gene expression association with learning using RNA-Seq, thereby focusing on transcriptional processes in memory formation. We also examined female behaviour (learning a novel spot preference) to examine if there was a temporal effect of differential gene expression and its associated learning behaviour. Our results showed that at the 1.5-hour time point, both learning and its associated differential gene expression was at its lowest, suggesting that there may be a correlational link between the lack of learning (mating outcome similar to naïve females) and the low transcriptional gene expression.

Future studies could explore this correlation in *B. anynana* or other systems. The only DEG that is shared between all five time points encoded for transcription initiation factor TFIID subunit 6 (*Taf6*), a subunit within a transcription complex TFIID, which plays an important role in initiating transcription by binding to the core promoter of the DNA [78,79]. Mutations in *Taf6* and other transcription associated factors (e.g. *Taf1, Taf13*) have been implicated in developmental disorders that affect intellectual disabilities in humans [80–85], suggesting that the disruption of normal TFIID function (through the mutations of its associated transcription factors) is detrimental to cognition. Further support for the importance of transcriptional processes to imprinting-like learning come from our WGCNA analyses, were the enriched GO terms for our significantly associated module with a similar significance temporal pattern to that of our DEG analysis, the lightgreen module, are also mostly related to translational and transcriptional processes. GO terms associated with transcriptional activity were also enriched in a study assessing associative learning in adult *C. elegans* [26], suggesting that the importance of transcriptional processes in long-term memory formation is conserved across the Ecdysozoa, and possibly learning context.

To further investigate whether molecular learning mechanisms are conserved across taxa and context, we compared our DEGs to those from various transcriptomic studies across different species, learning assays, and sensory modalities (Fig 4). Interestingly, *Hr38/ Nr4a* was found to be differentially expressed in multiple learning contexts and species, such as parasite-induced learning and rejection-induced courtship learning in *D. melanogaster* [49,51], auditory learning in female cowbirds[86], and aversive or spatial learning in mice [74–76,87] (Fig 4). Other differentially expressed genes in our data set, such as S-phase kinase-associated protein 2 (*BANY.1.2.g25897; skp2*) [45,49], have been implicated in cognition function and neuronal growth in Alzheimer’s studies using mice [88], as well as in eye development in *D. melanogaster* [89] (Fig 4). Paired box 2 (*BANY.1.2.g21950; Pax2/sv*), a member of the retinal determination gene network, is known to be primarily involved in eye/ photoreceptor development [90,91], but recent studies show that it also influences spatial learning in mice and autism spectrum disorders in humans and mice [92–95] (Fig 4).

**Fig 4.**
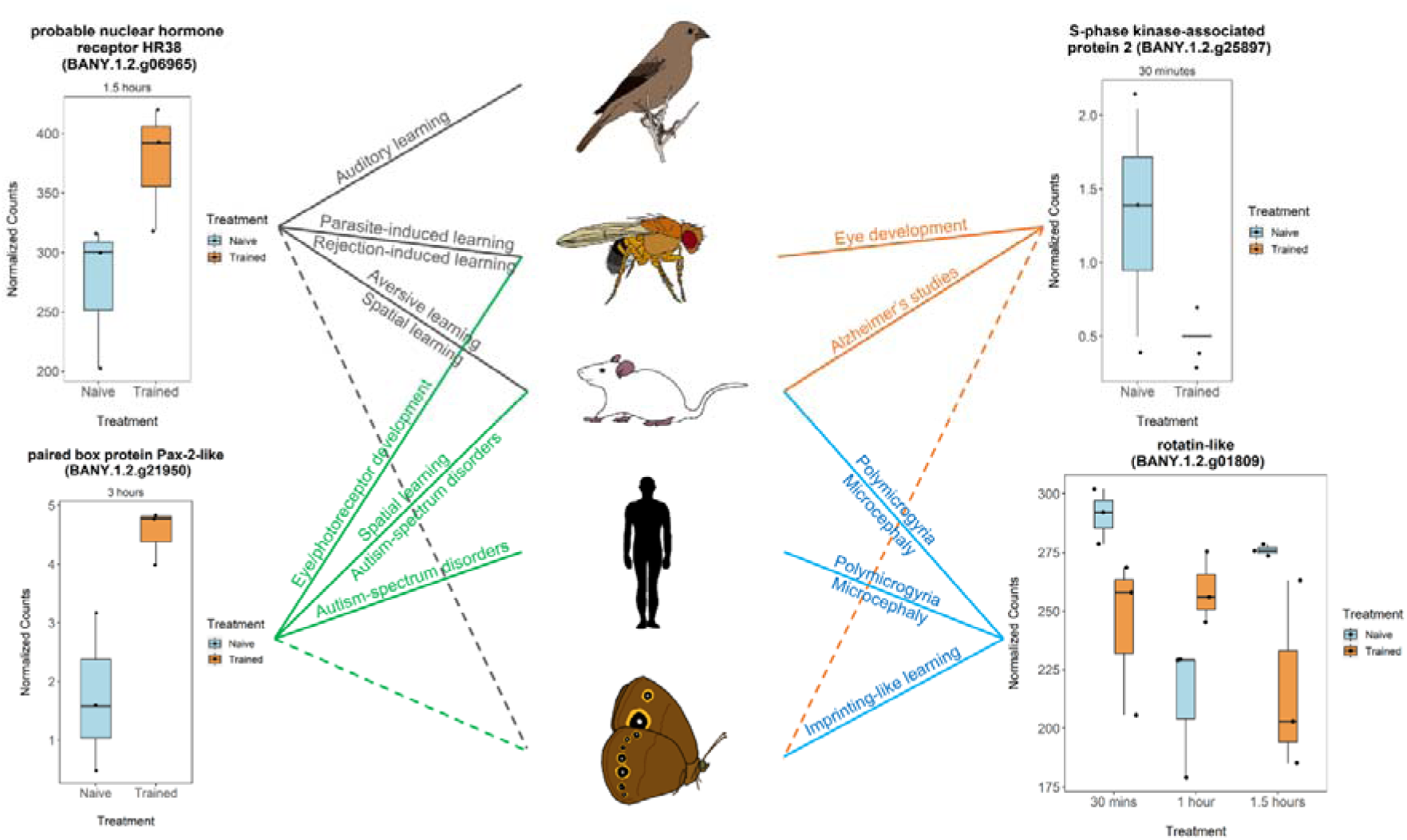
Putative learning and memory genes shared across taxa and learning contexts. Boxplots indicates the normalised counts of DEGs during the time point(s) when they are differentially expressed in our experiment, with full lines linking to the corresponding animal(s) and learning context. Dashed lines indicate that the corresponding DEG is also differentially expressed in our study. Blue indicates naive females, while orange indicates trained females.

Additionally, compared to a previous *B. anynana* study that investigated the gene expression networks behind sexually dimorphic imprinting-like learning [34], we found nine shared DEGs, including the rotatin-like protein (*RTTN/Ana3*), which is needed for the maintenance of sensory neurons [96,97]. *RTTN/Ana3* is also linked to human neurodegenerative diseases such as polymicrogyria and microcephaly, which can lead to learning disabilities in humans and mice [98–105] (Fig 4). Taken together, it would be worthwhile to functionally investigate whether these genes play a conserved role in learning and memory, especially since they are differentially expressed across multiple species (and sometimes phyla) and types of learning.

## Conclusions

This study shows the temporal dynamics of gene expression and mate preference learning behaviour in the butterfly *B. anynana*. We show that three hours of training had the strongest effect of learning compared to shorter time points, and there was no linear effect of training time on preference learning. For the first time, to our knowledge, we show that amount of differential gene expression does not always predict strength of behavioural response, as we found the highest DEG after one hour of exposure to a novel male spot phenotype, even though a one hour exposure does not induce a strong behavioural response in later mate choice assays. We also showed a decrease of learning and gene expression at the 1.5-hour time point, which is an intriguing correlation for future study. We also generated a list of putative learning and memory genes, some of which are cross-referenced with multiple transcriptomic studies across taxa and learning context, such as *Hr38*, *Pax2* and *Ana3*. Additional studies could test these genes for their functional role in mate preference learning. Lastly, our study supports the hypothesis that transcriptional processes play a major part in learning and memory formation across taxa and learning context.

## Materials and methods

### 1. Study Species and Animal Husbandry

*Bicyclus anynana* is a sub-Saharan, African butterfly that has been reared in the lab since 1988 [107]. The colony at the University of Arkansas was established in 2017, with ∼ 1,000 eggs from a colony at the National University of Singapore. *B. anynana* colonies at the University of Arkansas are reared in a climate controlled, USDA-APHIS approved (Permit # P526P-20-00417) greenhouse. The greenhouse was maintained at 27°C, with a relative humidity of 60-80%, and a 13:11 h light:dark photoperiod. The larvae were fed with maize plants (*Zea mays*) (Jollytime popcorn) *ad libitum* and adults were fed with rotten bananas. Adults that emerged on the same day (Day 0) were transferred to sex and age-specific cages. Since the laboratory colonies are large (∼1000), there is sufficient genetic diversity found in our colonies, and it is comparable to the genetic diversity found in natural populations [38,39].

### 2. Learning assays and the learning curve

To determine if the duration of exposure influences female ability to learn to prefer an unfamiliar, manipulated phenotype (four dorsal forewing UV reflective spots) over the innately preferred phenotype (two dorsal forewing UV reflective spots), we exposed newly emerged females to sexually mature 4-spotted males for different durations of time, and then tested subsequent female mate-choice. Exposure/training assays were performed one hour after sunrise. On date of emergence (Day 0), each female was paired with one Day 3, 4-spotted male for one of five different exposure times: i) 30 minutes, ii) 1 hour, iii) 1.5 hours, iv) 2 hours or v) 3 hours. We then removed the trainer males and confirmed that the trained females were unmated using a fluorescent powder (details below). The trained females were then kept isolated and fed for approximately 36 hours, allowing for long-term memory formation. We used naïve females (maintained in isolation since emergence) as control. For the mate-choice assays (Day 2), females were given a choice between 2 males: a 2-spotted male (innately preferred) and a 4-spotted male (innately unpreferred). To create these two male phenotypes, we manipulated male wing patterns using UV-reflective white paint with reflective spectra similar to the natural white eyespot centers [16]. For the 4-spotted butterflies, we painted two extra spots on the dorsal forewing surface. For the 2-spotted butterflies, we painted two spots directly on the natural dorsal forewing eyespots. Thus, male butterflies had different numbers of dorsal forewing UV reflective eyespots but the same amount of UV-reflective paint. All eyespot manipulations were done a day before behavioural (training or mate choice) assays.

We determined female mate choice by dusting the ends of females’ abdomens with fluorescent clownfish orange powder (Risk Reactor Inc; California, USA), and checking the abdomen ends of males using a blacklight flashlight [16]. The chosen male would have fluorescent orange powder residue on their abdomen ends. If both males had orange residue, or if no mating occurs after 48 hours, the mate choice assay was considered inconclusive and was not used. Note that this same method was used to confirm females did not mate during the training period. In the rare occasion when powder was transferred to the trainer male, that female was not used in future mate choice or gene expression assays.

### 3. Transcriptomic analysis of female whole heads across time points

#### 3.3. Sample collection, RNA extraction and library preparation

To determine the gene expression changes across time, we extracted RNA and prepared RNA libraries from female whole heads for RNA-Seq by decapitating three naïve (socially isolated) and three trained females with RNA-free scissors at each time point. Each head was placed into individual RNA-free 1.5ml LoBind tubes and immediately flash frozen in liquid nitrogen. Frozen samples were then stored in −80LJC until dissection. To prevent RNA degradation during dissection, we soaked the heads in 500 ul of prechilled RNAlater ICE (Ambion; Texas, USA) and kept them in −20LJC at least 16 hours before dissection and RNA extraction. We then dissected the heads in RNAlater ICE, and extracted RNA using the Nucleospin miRNA kit (Macherey-Nagel; Duren, Germany), which extracts both large RNA (>200 bp) and small RNA (<200 bp). We utilized large RNA for downstream library preparation and analyses. Quality of RNA were determined using Nanodrop 2000 (Thermo Fisher Scientific; Massachusetts, USA) and TapeStation 2200 (Agilent; California, USA). We prepared RNA libraries using the KAPA mRNA HyperPrep Kit and Unique Dual-Indexed Adaptors (KAPA Biosystems; Massachusetts, USA), with 200 ng of RNA as input, and shipped to the University of Chicago’s Genomics Facility on dry ice. The libraries were further accessed for quality using a 5300 Fragment Analyser (Agilent; California, USA), followed by a 50 bp single-end sequencing across four lanes of NovaSeq 6000 (Illumina; California, USA).

#### 3.2 Differential gene expression analysis

We concatenated the raw fastqc files from each library and accessed their quality using FASTQC v0.11.5, removed adaptor sequences using Trimmomatic v0.38, and aligned the trimmed sequences to the most recent *B. anynana* reference genome (v1.2)[60] using STAR v2.7.1a with the default parameters and the “—twopassMode Basic” option, and quantified using HTSeq.

We then performed permutation tests by generating 1000 permutated datasets by randomly assigning each sample name for the entire dataset in R [40,41] (N= 30). We then performed GLM in the same way as the actual data, generating a null distribution of 1000 permutated p-values per gene transcript, and a gene was considered differentially expressed when its associated p-value fell below the 1% tail of the permutated data p-value distribution. This method has been shown to better capture the structure of data and does not assume gene independence [42]. Therefore, this reduces the risk of overcorrection compared to other multiple test correction methods, since gene expression is likely influenced by other associated genes, and thus not independent. Pairwise comparisons of interest (e.g. Naïve 30 minutes compared to Trained 30 minutes, to compare effect of training in specific time point) were extracted and analysed using the “DESeq2” package [43].

We used the GLM “y ∼ group”, where group combines time point and training status (e.g. Naïve 30 mins, Trained 30 mins, Naïve 1 hour…; for a total of 10 different sample groups). This design allowed us to contrast the effect of training at each time point. Genes with a total read alignment count of <10 were filtered out and not included in the differential expression analysis. Gene expression was calculated as the log of the expression fold change (log_2_FC) and the “apeglm” method [44] was used for log_2_FC shrinkage. Gene ontology (GO) term annotations for all differentially expressed genes were extracted from the *B. anynana* reference genome functional annotation from Ernst & Westerman (2021). We then collected the amino acid sequences of the differentially expressed genes (DEGs) from LepBase[46], and used BLASTP [47] to search the NCBI database for orthologs. The top candidate ortholog was determined based on the lowest E-value. In cases where numerous hits had identical E-values, we selected based on the highest hit score. If there were no hits using BLASTP, we then queried the gene transcript from Lepbase and used BLASTN instead. If there were no hits using either BLASTP and BLASTN, we then defined the gene as NA.

#### 3.2 Identification of gene categories

To categorise genes into their different putative functions, we used four different methods. Firstly, for putative vision and chemosensory (gustatory, olfactory) genes, we cross-referenced the gene list from [45]. Secondly, we then manually searched for gene orthologs in *Drosophila melanogaster* using FlyBase v FB2022_03 [48], since most of BLASTP’s hits were gene names from humans or other mammalian model species, and grouped the genes based on their GO terms in Flybase. Thirdly, we cross-referenced published DEGs associated with learning and neurodevelopment from other peer-reviewed transcriptomic studies [40,49–53]. Lastly, we used NCBI’s Gene Expression Omnibus Profiles (GEO)[54] database to query our differentially expressed genes, specifically searching for studies pertaining to learning and memory. We manually searched for terms like “learning”, “memory”, “brain”, and “mushroom body” alongside our DEGs. Four gene expression datasets [55–58] were chosen since they showed up repeatedly throughout our searches, either indicating that their datasets were relevant to our study and/or that the datasets were large. Using GEO’s embedded GEO2R program, we analysed the two of the four chosen datasets [57,58] individually because there were no published gene lists available for them. This was done by comparing gene counts between their “control” and “mutant” groups, and we retrieved two lists of genes that were differentially expressed at p < 0.05.

We categorized all genes into one of eight categories based on the GO terms: “learning and memory”, “visual genes”, “chemosensory genes”, “nervous system”, “hormones”, “ion channel”, “uncharacterised” and “other”. For “learning and memory”, genes under this group were implicated as candidate genes in either published transcriptomic studies[40,49–52,55], two GEO datasets[57,58] or by GO terms of their orthologs in Flybase. This included any GO terms associated with learning and memory (e.g. long term memory, short term memory, olfactory memory, visual learning, associative memory). “Visual genes” and “chemosensory genes” were genes from Ernst & Westerman (2021) and GO terms in Flybase. This included any GO terms involved with vision, smell or sound (e.g., phototransduction, compound eye photoreceptor development, rhabdomere development/ morphogenesis, sensory perception of sound/ smell/ mechanical stimulus, antennal development, response to stimulus). For “nervous system”, genes included were found in two published transcriptomic studies[53,56] or GO terms in Flybase (e.g. neurogenesis, regulation of nervous system development, axon/ dendrite morphogenesis, axon/ dendrite guidance, axonal/ dendritic transport, synaptic growth/ assembly/ organisation/ transmission, synaptic growth at muscular joint, neurotransmitter release/ synthesis). For “hormones”, this included genes with GO terms involved in the regulation or synthesis of hormones, as well as the hormones themselves. For “ion channel”, this included genes with GO terms encoding for ion channels, or subunits of an ion channel. For “uncharacterised”, this included genes that are expressed in the open reading frame but there was no experimental evidence of their translation. Lastly, for “other”, this included all genes that do not fall under any of the aforementioned groups.

#### 3.3. Weighted gene co-expression network analysis

To identify correlation patterns between genes across different time points, we performed weighted gene co-expression network analysis (WGCNA) using the “WGCNA” package in R [59]. The data was pre-processed by filtering out genes with reads <10 to minimise noise from lowly expressed genes. We then performed variance-stabilising transformation on the data using “varianceStabilizingTransformation” function in the “DESeq2” package. We also manually created binary indicators for our “traits” (i.e., under the “training” variable, we defined “0” for naïve and “1” for trained individuals; under “time” variable, we defined “0” for the 30 min time point, “1” for the 1 hour time point, “2” for the 1.5 hour time point, “3” for the 2 hour time point, “4” for the 3 hour time point). Then, we used the “binarizeCategoricalVariable” function to create pairwise contrasts for all possible permutations, and filter out our pairwise contrasts of interest (i.e. naïve 30 mins vs trained 30 mins, naïve 1 hour vs trained 1 hour, naïve 1.5 hours vs trained 1.5 hours, naïve 2 hours vs trained 2 hours, naïve 3 hours vs trained 3 hours, time, and training). An adjacency matrix with type = “signed”, and a topological overlap matrix (TOM) with TOMType = “signed” were constructed with the softPower threshold set to 8. We then identified modules of co-expressed genes using the “cutreeDynamic” function of parameters deepSplit = 2, pamRespectsDendro = FALSE and minClusterSize = 30. After that, we merged modules of high co-expression similarity by calculating and clustering their eigengenes in a dendrogram with cutHeight = 0.25. A module-trait heat map was then created by comparing the association between a specific module and “trait” (R^2^ value), adjusting all p-values using the FDR method, with any FDR < 0.05 considered as significant.

### 4. Behavioural statistical analysis

We recorded the mating outcomes of females with 2- or 4-spotted males during the mate choice assays. For analysing proportion of females mating with 2- or 4-spotted males, we used a chi-square analysis comparing different time points to the naïve control (n = 35). All analyses were done in R.

### 5. Ethical Statement

All *B. anynana* butterflies are maintained in laboratory conditions specified by U.S. Department of Agriculture APHIS permit (# P526P-20-00417). Butterflies were fed with *ad libitum* food and water. Those used in this experiment were decapitated quickly and flash frozen immediately after for this and future studies.

## Data availability statement

All raw sequence data associated with this study have been made accessible through the NCBI Sequence Read Archive (SRA) database under BioProject PRJNA1030143. Behavioural data and codes are available at Dryad (doi: 10.5061/dryad.wstqjq2tz).

## Acknowledgements

We would like to thank Grace Hirzel, Matthew Murphy, Sushant Potdar, Kiana Kasmaii, Chance Powell, Jonas Amenyo, Keity Farfan-Pira and Dimitry Kutcherov for their assistance with animal husbandry. This research was supported by NSF IOS-1937201 to ELW and the University of Arkansas. We would also like to acknowledge the Arkansas High-Performance Computing Centre, which is funded through multiple National Science Foundation grants and the Arkansas Economic Development Commission.

---

**Fig S1. Gene groupings across five time points.** “Learning and memory” was denoted in green, “Vision” in blue, “Chemosensory” in red, and “Nervous system” in yellow. “Hormones”, “Ion channel”, “Uncharacterised” and “Other” are not represented here.

**Fig S2. Hr38 expression across five time points.** “Naïve” females are denoted in blue, while “Trained” females are denoted in orange. There is significant difference in Hr38 expression at the 1.5-hour time point, but not at other time points.

**Fig S3. GO enrichment plots for the lightgreen module.** (A) The original GO enrichment plot with all significantly enriched GO terms. (Bs) The reduced GO enrichment plot with the most specific GO terms. For each GO term, the percentage of sequences annotated with that GO term within the lightgreen module is plotted along with the percentage of sequences annotated with that same GO term in the Reference set (i.e. all genes expressed).

