## Supplemental Figures S1-3 for "Asynchronous temporal variance in learning behaviour and neural gene expression in a butterfly"

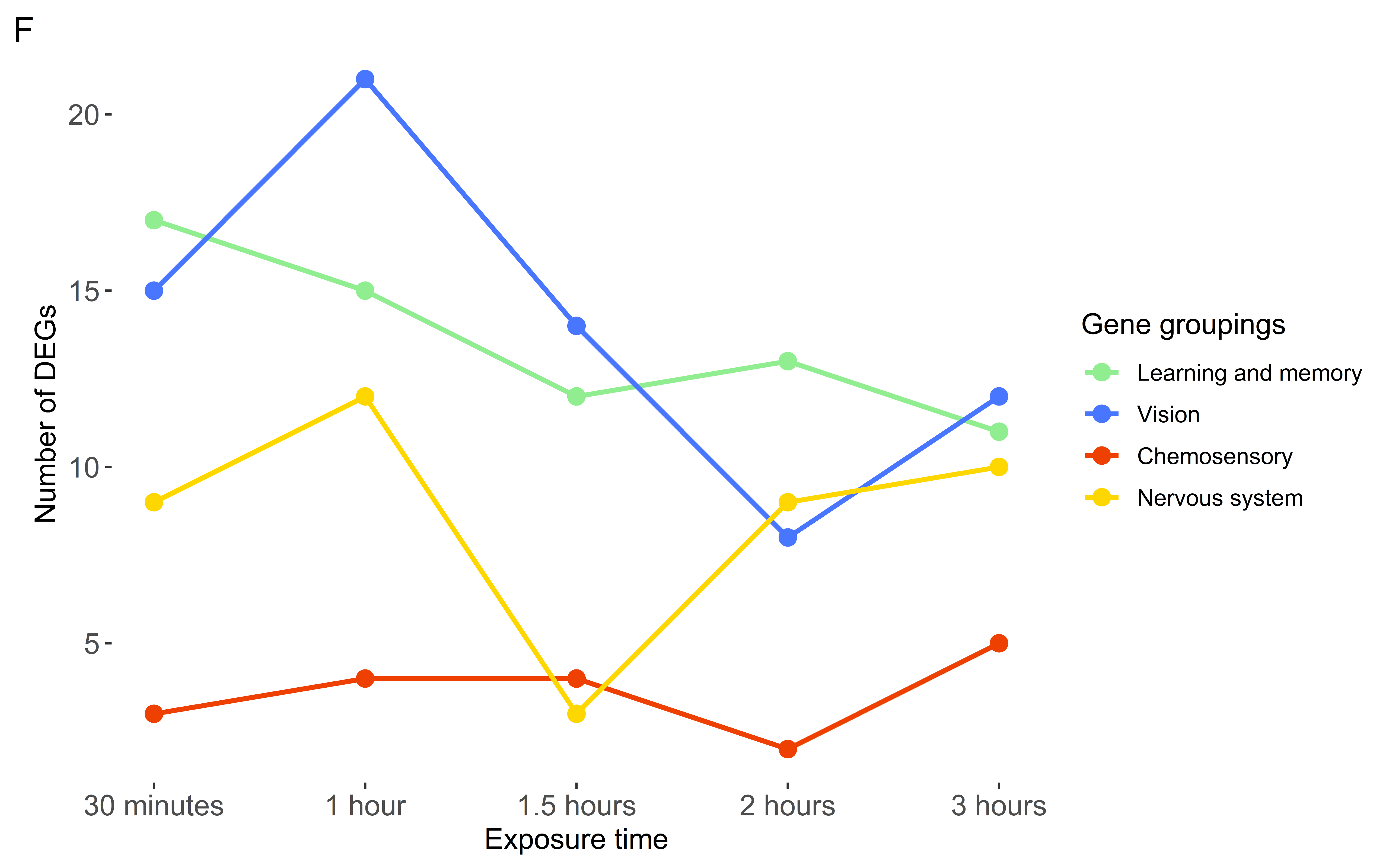

**Figure S1. Gene groupings across five time points.** “Learning and memory” was denoted in green, “Vision” in blue, “Chemosensory” in red, and “Nervous system” in yellow. “Hormones”, “Ion channel”, “Uncharacterised” and “Other” are not represented here.

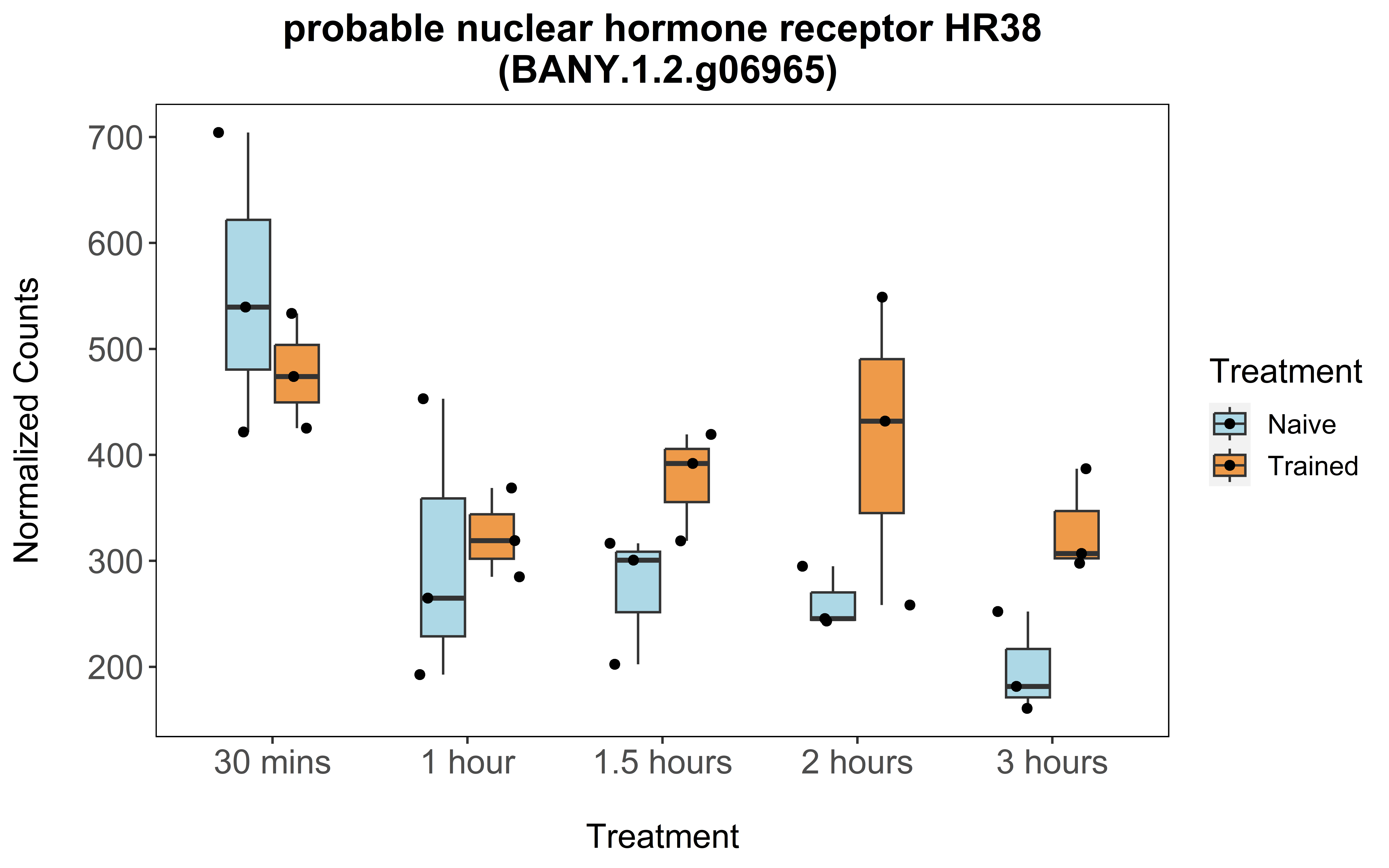

**Figure S2.** Hr38 expression across five time points. “Naïve” females are denoted in blue, while “Trained” females are denoted in orange. There is significant difference in Hr38 expression at the 1.5-hour time point, but not at other time points.

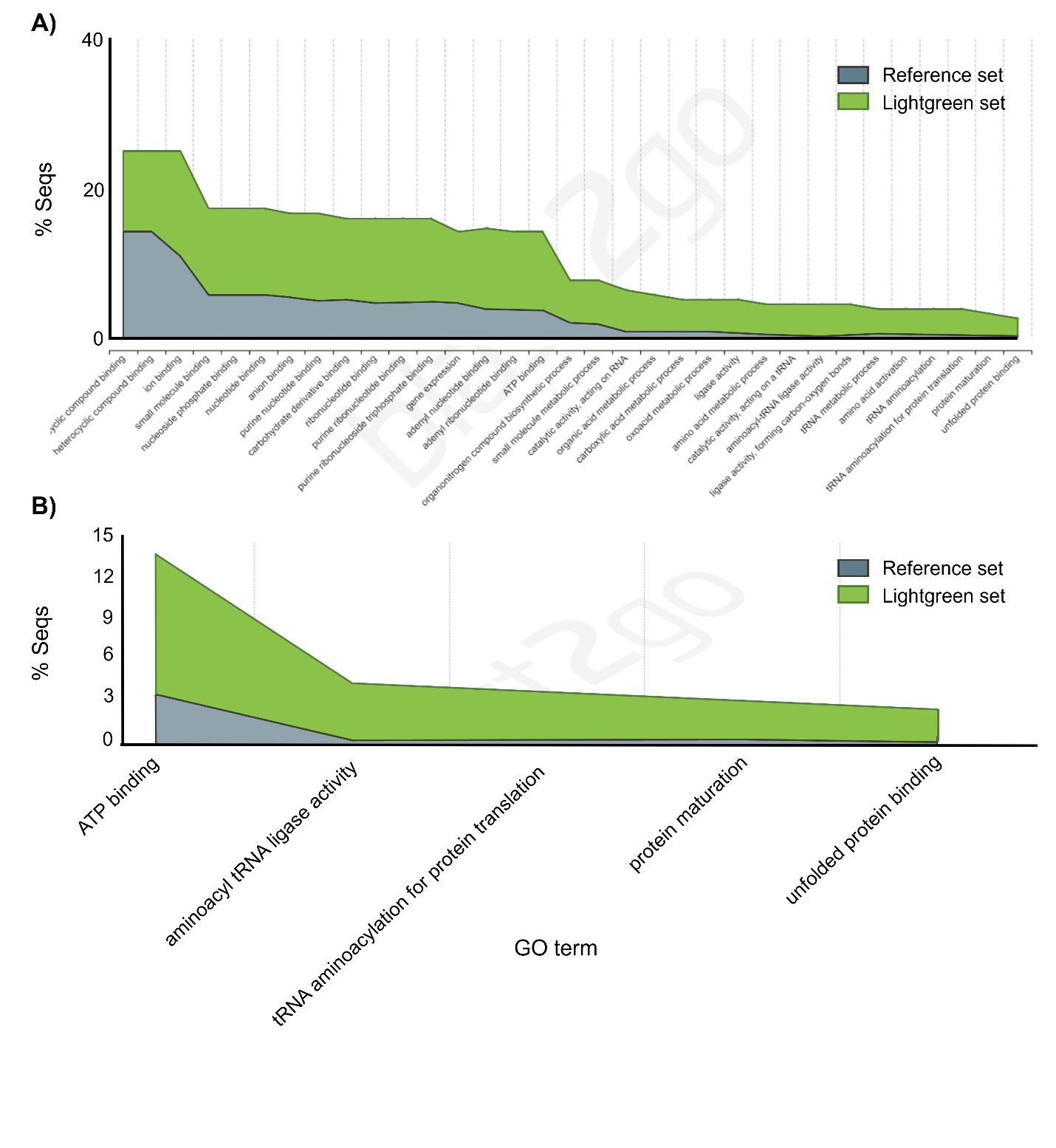

**Figure S3.** GO enrichment plots for the lightgreen module. (a) The original GO enrichment plot with all significantly enriched GO terms. (b) The reduced GO enrichment plot with the most specific GO terms. For each GO term, the percentage of sequences annotated with that GO term within the lightgreen module is plotted along with the percentage of sequences annotated with that same GO term in the Reference set (i.e. all genes expressed).
